# Radial Glia cell infection by *Toxoplasma gondii* disrupts brain microvascular endothelial cell integrity

**DOI:** 10.1101/378588

**Authors:** Daniel Adesse, Anne Caroline Marcos, Michele Siqueira, Cynthia M. Cascabulho, Mariana C. Waghabi, Helene S. Barbosa, Joice Stipursky

## Abstract

Congenital toxoplasmosis is a parasitic disease that occurs due vertical transmission of the protozoan *Toxoplasma gondii* (*T. gondii*) during pregnancy. The parasite crosses the placental barrier and reaches the developing brain, infecting progenitor, glial, neuronal and vascular cell types. Although the role of Radial glia (RG) neural stem cells in the development of the brain vasculature has been recently investigated, the impact of *T. gondii* infection in these events is not yet understood. Herein, we studied the role of *T. gondii* infection on RG cell function and its interaction with endothelial cells. By infecting isolated RG cells with *T. gondii* tachyzoites, we observed reduced cell proliferation and neurogenesis without affecting gliogenesis levels. Conditioned medium (CM) from RG control cultures increased ZO-1 and β-catenin protein levels and organization on endothelial bEnd.3 cells membranes, which was completely impaired by CM from infected RG, resulting in decreased transendothelial electrical resistance (TEER). Cytokine Bead Array and ELISA assays revealed the presence of increased levels of the pro-inflammatory cytokine IL-6 and reduced levels of anti-inflammatory cytokine TGF-β1 in CM from *T.gondii*-infected RG cells. Treatment with recombinant TGF-β1 concomitantly with CM from infected RG cultures led to restoration of ZO-1 staining in bEnd.3 cells. Our results suggest that infection of RG cells by *T. gondii* modulate cytokine secretion, which might contribute to endothelial loss of barrier properties, thus leading to impairment of neurovascular interaction establishment.

## 1) INTRODUCTION

Toxoplasmosis is a parasitic disease that affects all warm-blooded animals, including humans. The disease is caused by a protozoan parasite, *T. gondii* and has a high global seroprevalence, estimated in approximately 1/3 of the world’s population (Dubey, 2010). Transmission occurs by ingestion of uncooked meat from infected animals, that contains tissue cysts, or by ingestion or inhalation of sporulated oocysts, shed with feces of infected felids. The cysts are digested by proteolytic enzymes present in the stomach and small intestine, which then release infective parasites that rapidly invade epithelial cells of the small intestine and differentiate into fast-replicating tachyzoite forms. After intense intracellular proliferation, parasites promote host cell lysis and can disseminate throughout the entire organism (reviewed in Hill and Dubey, 2016). During the acute phase, patients may present lymphadenopathy, which may be associated with fever, fatigue, muscle pain, sore throat and headaches (Montoya and Liesenfeld, 2004).

*T. gondii* can also be vertically transmitted during gestation, leading to Congenital Toxoplasmosis (CT), established by the capacity of the parasite to cross the placental barrier and reach the developing brain tissue, where both tachyzoites and tissue cysts can be found in the developing brain parenchyma (Ferguson et al., 2013). CT is part of the TORCH complex of infectious diseases (**<u>T</u>**oxoplasma, **<u>R</u>**ubella, **<u>C</u>**ytomegalovirus, **<u>H</u>**erpes simplex 2. **<u>O</u>** stands for Others, and includes chlamydia, HIV, Coxsackievirus, Syphilis, Hepatitis B, chicken pox and Zika virus) that can be transmitted from the mother to the fetus (Neu et al., 2015; Mehrjardi, 2017). Infection with some of these pathogens can potentially lead to congenital defects including, but not limited to, microcephaly, growth and mental retardation, heart disease and hearing loss (Neu et al., 2015; Klase et al., 2016). Although transmission during the third trimester has been implicated in reduced impact on the fetus, infection during the first trimester is extremely disruptive, with severe neurological manifestations including microcephaly, cognitive/intellectual disabilities, deafness and blindness (Wallon et al., 1999).

Deleterious effects of infection of mouse neural progenitor cells by a highly infective *T. gondii* strain was linked to apoptosis induction by endoplasmic reticulum stress signaling pathway activation (Wang et al., 2014). In addition, reduced neuron and astrocyte generation from the neural C17.2 stem cell line by disruption of the Wnt/β–catenin signaling pathway have also been suggested as an underlying mechanism of *T. gondii*-induced neural pathological damage during brain development (Gan et al., 2016; Zhang et al., 2017).

RG cells are the major multipotent neural stem cell population present during the embryonic cerebral cortex development period and originate most of the neuronal and glial cell types found in neural tissue, by activation of multiple signaling pathways (Gotz and Barde, 2005; Kriegstein and Alvarez-Buylla, 2009; Stipursky et al., 2012; Stipursky et al., 2014). Besides its well-known role as neural stem cells, RG have recently been demonstrated to directly control vascular development and blood brain barrier (BBB) formation in the embryonic cerebral cortex (Ma et al., 2013; Errede et al., 2014; Hirota et al., 2015; Siqueira et al., 2017).

We have previously demonstrated that the gliogenic and neurogenic potential of RG cells during cerebral cortex development is controlled by TGF-β1 signaling pathway activation, both *in vitro* and *in vivo* (Stipursky et al., 2012; Stipursky et al., 2014). Neuroepithelial and RG neural progenitors interact with immature endothelial cells, derived from the perineural vascular plexus (PNVP) that surrounds the neural tissue early during the embryonic period. Such an interaction is essential to promote invasion of endothelial cells and vascularization of the developing CNS. Endothelial cells from the PNVP invade forebrain tissue as early as E9.5 in mice and migrate towards the ventricular surface, guided by VEGF gradients secreted by neural progenitor cells (Bautch and James, 2009; Anderson et al., 2011; Liebner et al., 2011; Takahashi et al., 2015). Recently, we demonstrated that RG cells coordinate the formation of the vascular tree of the brain, by controlling angiogenesis in the developing cortex. Specifically, RG cells secrete a vast repertoire of pro-angiogenic factors, including TGF-β1 and VEGF-A, that induce endothelial proangiogenic genes expression and regulate migration and blood vessel branching in the embryonic cerebral cortex (Siqueira et al., 2017).

Blood vessel development and neural cell generation in the CNS are essential steps for the establishment of the BBB. The BBB is a multicellular structure formed by capillary endothelial cells, astrocytic endfeet, pericytes and neighboring microglia and neurons, that control the transport of nutrients, oxygen and other substances, and prevent the free passage of toxic agents and pathogens (Kim et al., 2006; Anderson et al., 2011). In the adult brain, *T. gondii* can infect endothelial cells (Konradt et al., 2016) and, by modulation of adhesion proteins, contribute to decreased adhesion between adjacent endothelial, allowing for transmigration of inflammatory cells into the brain parenchyma (Lachenmaier et al., 2011).

Although RG physiology greatly determines the correct formation of the cerebral cortex, including its vascularization (Ma et al., 2013; Errede et al., 2014; Hirota et al., 2015; Siqueira et al., 2017), the understanding of the impact of *T. gondii* infection on RG-endothelial interactions in the embryonic CNS has never been addressed.

Here, we investigated the role of *T. gondii* infection on RG physiology and its potential to control endothelial barrier properties establishment. We demonstrated that infection reduce the neurogenesis potential and altered the RG secretome. Such altered alterations lead to important dysfunctions in microvascular brain endothelial cells, presenting reduced tight junction stability and barrier properties when incubated with a conditioned medium obtained from infected RGs.

## 2) MATERIAL AND METHODS

### 2.1) Compliance with Ethical Standards

All animal protocols were approved by the Federal University of Rio de Janeiro Animal Research Committee (CEUA 041/14). Animals were housed in a temperature-controlled room with a 12/12 h light/dark cycle and allowed food and water *ad libitum*.

### 2.2) Toxoplasma gondii infection

Parasites from the ME49 strain were obtained from the brains of C57/bl6 mice infected 45 days before isolation. Cysts were ruptured with an acid pepsin solution and free parasites were added to monolayers of Vero cells (ATCC^®^ CCL-81™). After two weeks of culture re-infections, tachyzoites released from the supernatant were collected and centrifuged. Cell cultures were infected with the tachyzoites at a multiplicity of infection (MOI) of 3 parasites:host cell (Luder et al., 1999) for 2 hours in 300 μL of DMEM-/F12/2% B27/ Penicilin-streptocycin (Thermo) (RG cultures). Cells were washed in Ringer’s solution, fresh culture media was added, and cultures were kept for additional 22 hours, when conditioned medium was collected and cells fixed for immunostaining. Mock-infected cultures (treated with fresh culture medium) were used as controls.

### 2.3) Radial glia (RG) cultivation

RG isolation from E14 gestational day-old Swiss mouse embryos was carried out as previously described by Stipursky et al. (2014). Briefly, gestational day14 Swiss mice embryos were collected and dissected for cerebral cortex separation. After dissection, tissues were dissociated in DMEM/F12 Glutamax high glucose (Thermo) medium and after cell counting, 3.10^5^cells were plated in 25 cm2 culture flasks in neurosphere “growing media” DMEM/F12 Glutamax high glucose (Thermo) containing 0.1% penicillin/streptomycin, 2% B27 (Thermo), 20ng/mL EGF (epidermal growth factor, Thermo) and 20 ng/mL FGFb (basic fibroblast growth factor, R&D Systems), for 6 days, in *vitro*. 2/3 of the media was changed every 2 days. After this period, neurospheres were enzymatically dissociated in 0.05%Trypsin/EDTA (Thermo). After cells isolation, 2.10^5^ cells were plated on glass coverslips previously coated with 5 μg/mL laminin (Thermo) and incubated in DMEM/F12 Glutamax (high glucose) without serum and supplemented with 2% B27, 20 ng/mL FGFb and EGF (Thermo). Twenty-four hours after plating, cells were infected with the tachyzoite forms of the *T. gondii*, ME49 strain as described above. After infection, cells were gently washed to remove extracellular parasites and 300 μL of fresh DMEM/F12/ medium were added, followed by 22 hours of incubation. Next, non-infected and infected RG cells were fixed with paraformaldehyde 4% solution in PBS (Sigma-Aldrich) for immunocytochemistry assays. Supernatants were collected, centrifuged for 10 min at 10,000 rpm (4 °C) to eliminate cell debris and extracellular parasites and frozen at − 80 °C to be further used as a Conditioned Medium (CM) or for cytokine measurements.

### 2.4) Enzyme-Linked Immunosorbent Assay (ELISA)

TGF-ß1 levels present in the conditioned medium derived from non-infected (RG-CM) and infected (Inf-RG-CM) RG cells, were measured by the Mouse TGF-ß1 ELISA DuoSet Kit (R&D Systems) following the manufacturer’s instructions.

### 2.5) Cytometric Bead Array (CBA)

Cytokine levels were evaluated by flow cytometry in culture RG-CM and Inf-RG-CM supernatants. IL-10, IL-17, TNF, IFN-γ, IL-6, IL-4 and IL-2 were detected using a Cytometric Bead Array (CBA) Th1/Th2/TH17 kit (BD), according to the manufacturer’s instructions. Samples were acquired using a FACScalibur flow cytometer (BD), and the data analysis was performed using the CBA analysis FCAP software (BD).

### 2.6) bEnd.3 cell line cultivation

A total of 6.10^4^ murine brain microvascular endothelial cells (bEnd.3, ATCC^®^ CRL-2299™) were plated on glass coverslips previously coated with 0.01% porcine gelatin solution (Sigma-Aldrich) in bEnd.3 medium [DMEM/F12 Glutamax high glucose (4500 mg/L) with 10% heat-inactivated Fetal Bovine Serum (Cultilab) and 1% penicillin/streptomycin solution (Thermo)] for 14 days, with the medium changed every 2 days. After reaching confluence, cultures were treated with RG-CM, Inf-RG-CM, Inf-RG-CM+TGF-β1 (10ng/Ml, R&D Systems) or TGF-β1 (10ng/mL) for 24 hours. Cultures were fixed with PFA 4% for immunocytochemistry assays. Cells were used between passages 25 to 30.

### 2.7) Immunocytochemistry

Immunostaining was performed as previously described by Siqueira et al. (2017). Briefly, fixed cultures were permeabilized for 5 min with 0.05% Triton x-100 solution in PBS and non-specific binding blocked by incubation with blocking solution containing 5% Bovine Serum Albumin (BSA - Sigma-Aldrich)/2.5% Normal Goat Serum (NGS)/PBS for 1 hour. Cells were incubated with primary antibodies, diluted in blocking solution and maintained overnight at 4°C. For RG cultures, immunostaining primary antibodies were: mouse anti-Nestin (marker of neural progenitor cells, Millipore, 1:200); rabbit anti-BLBP (marker of radial glia cells, Chemicon, 1:500), rabbit anti-Ki67 (nuclear marker of mitotic cells, Abcam, 1:100), rabbit anti-GFAP (intermediary filament protein specific to glial cells, Dako Cytomation, 1:500) and mouse anti-β-III-tubulin (specific isoform found in immature neurons, Promega, 1:1000), and mouse anti-cleaved Caspase 3 (apoptosis cell marker, Abcam, 1:100). For endothelial culture immunostaining, primary antibodies were: mouse anti-ZO-1 (Invitrogen, 1:300), rabbit anti-β-catenin (Sigma-Aldrich, 1:200). Subsequently, cells were extensively washed in PBS and incubated with secondary antibodies, conjugated to AlexaFluor 488 or AlexaFluor 546 (Thermo), for 2 h at room temperature. Nuclei were DAPI-labeled (4’, 6-Diamidino-2-phenylindole; Sigma-Aldrich). Glass coverslips were mounted in glass slides with Faramount mounting media (Dako Cytomation) and visualized under a fluorescence optical microscope Nikon TE3000 or a Leica SPE confocal microscope. Fifteen random images under a 40x objective were acquired from each glass coverslip from at least 3 independent experiments done in triplicate.

### 2.8) Trans-Endothelial Electrical Resistance (TEER)

bEnd.3 cells were plated onto 0.01% gelatin-coated Transwell inserts (Falcon) with 3 μm pores at a density of 10^5^ cells per insert. Cultures were maintained in bEnd.3 medium at 37°C in 5% CO_2_ atmosphere and resistance was measured daily using a Millicell-Electrical Resistance System (Millipore, Bedford, MA) with an adjustable electrode (“chopstick electrode^, MERSSTX03), as described by Srinivasan et al., 2015. Cells reached confluence after approximately 14 days (minimum of 60 Ω × cm^2^), with the medium changed every 2 days. One electrode is inserted into the upper trans-well insert compartment and the other electrode to the lower compartment. Care is taken to ensure that all compartments have the same volume of medium across biological and technical replicas (300 ml in the upper compartment and 600 ml in the lower). A square wave current of 12.5 Hz is applied to the electrodes and the resulting current is measured. To calculate TEER, the background resistance reading from an empty insert was subtracted from the resistance reading for each condition and the result was multiplied by 0.33, relative to the insert area, and results were expressed as Ω × cm^2^. One insert per experiment was maintained in bEnd.3 medium (10% FBS), while experimental data was obtained from cultures incubated with DMEM/F12 high glucose with antibiotics solution and no FBS. Cells were used for experiments when TEER reached a minimum of 60 Ω × cm^2^. TEER was obtained before experimental procedures (t=0) and 24 h after treatments with CMs or infection (t=24). The variation index for each experimental condition was calculated as TEER_t=24_/TEER_t=0_.

### 2.9) Quantification and statistical analyses

Quantification analyses of cell populations (RG, neurons and astrocytes) were carried out manually using the Photoshop CS6 software. The percentage of each stained cell population, in each microscopic field, was calculated in relation to total DAPI stained cells numbers in the same field. bEnd.3 labeling intensity analysis was carried out using the ImageJ software and the TiJOR analysis was performed using the TiJOR macro for ImageJ, which in an index of localization of tight junction proteins in membrane-membrane contact region of adjacent cells as described by Terryn et al. (2013). The GraphPad Prism 6.0 software was used for the statistical analyses, obtained at http://www.graphpad.com/scientific-software/prism. Statistical significance from at least 3 independent experiments was determined by unpaired t-test and ANOVA for biological effects with an assumed normal distribution. P value <0.05 was considered statistically significant.

## 3) RESULTS

### 3.1) *T. gondii* infection alters Radial Glia cell numbers and neurogenesis

In order to understand the effects of *T. gondii* infection on the RG differentiation potential, isolated RG cells from E14 cerebral cortex were infected with the tachyzoite forms of the parasite. We used the cells 24 hours after neurosphere dissociation and plating. At this time, Radial glia cultures are virtually pure with 96% of cells positive for blbp and 95.7% positive for nestin (92% were blbp/nestin double positive, Stipursky J., personal communications). After 24 hours of infection with *T. gondii*, mock-infected (control) cells displayed typical radial bipolar morphology and expression of typical RG neural stem cells markers BLBP and Nestin, *in vitro* (**Figure 1A**). Parasites were detected in the cytoplasm of Nestin-positive cells (**Supplementary Figure 1C-D**). Infected cultures presented a 40% decrease in the number of BLBP/Nestin double-labeled cells (**Figure 1 A-C**). This event was accompanied by a 25 % decrease in proliferative RG cells double labeled for Ki67 and Nestin markers (**Figure 1 D-E**). In parallel, *T. gondii* infection also significantly decreased the numbers of early neurons (as detected by β-III-tubulin labeling) by 36% (**Figure 1 G-I**). However, no effect on the number of astrocytic (GFAP-positive) cells was detected in infected cultures compared to the controls (Figure 1 J-L). The average of total cells number of β-TubulinIII and Nestin positive cells double labeled for the apoptotic marker cleaved-Caspase 3 was not affected by infection (**Supplementary Figure 2A**). Overall, *T. gondii*-infected cultures presented a shift in the percentage of cell composition when compared to uninfected controls, with Nestin/BLBP and β-III-tubulin positive cells being directly affected, although other unidentified cell types also tended to present percentage alterations, increasing from 17% in controls to 40% in infected cultures (**Figure 1M**).

**Figure 1.**
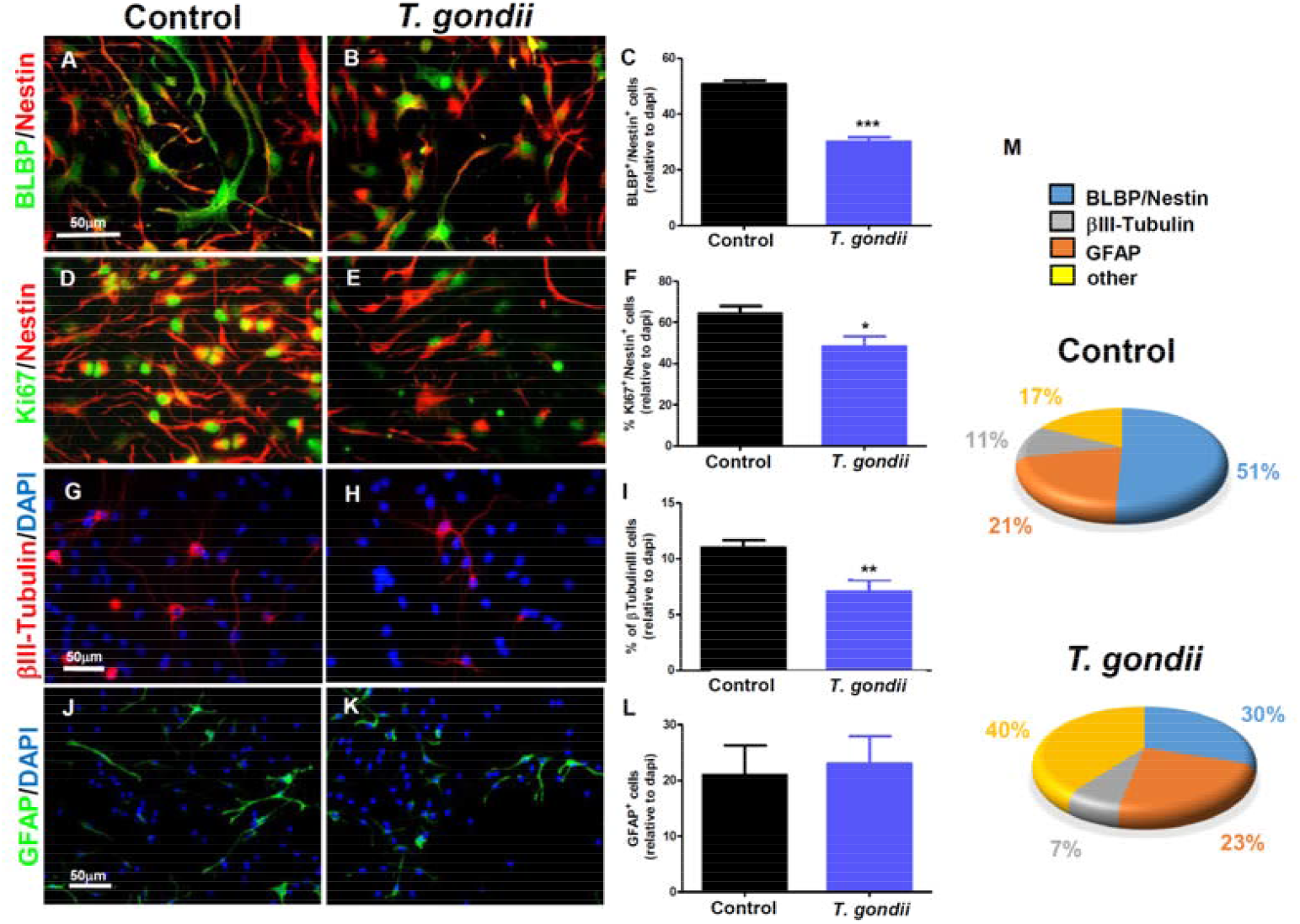
*T. gondii* infection decreases Radial glia cell proliferation and neurogenic potential. Infection of RG cells cultures with *T. gondii* tachyzoites for 24 h significantly decreased the number of Nestin/BLBP double labeled cells number, compared with control (A-C) which was accompanied by reduced numbers of proliferative Ki67/Nestin positive cells compared with control uninfected cells (D-F). A significant decrease in generation of β-III-tubulin positive neuron numbers was induced by *T. gondii* infection, when compared with control (G-I). RG differentiation into GFAP positive astrocytes was not affected by *T. gondii* infection when compared to controls (J-L). *T. gondii* infection induced significant changes in the percentage of cell composition in RG cultures compared to controls (M). *P=0.0423, **P=0.0019, ***P=0.0001 versus controls, unpaired *t* test.

### 3.2) *T. gondii* infection affects RG potential to control endothelial barrier property

To investigate the potential of RG cells to control endothelial cells function and the role of *T. gondii* infection in this context, we cultivated b.End3 endothelial cells until confluence. Through immunocytochemistry analyses, we identified ZO-1 adapter tight junction protein mainly distributed along cell-cell contacts in the control condition (**Figure 2A**). Treatment of endothelial cells with conditioned medium derived from uninfected RG cells (RG-CM) for 24 h significantly increased the ZO-1 labeling intensity levels by 36% (**Figure 2B, D**). This was concomitant with an increase in organization levels of the tight junctions (TiJOR) by 90% (**Figure 2 E**). However, treatment of endothelial cells with conditioned medium derived from infected RG cells (Inf-RG-CM) completely abrogated RG potential to induce ZO-1 labeling intensity, by 40%, when compared to control cultures, although the TiJOR index did not differ from the controls (**Figure 2 C, D**). In summary, Inf-RG-CM-treated presented deleterious effects on tight junction protein levels and organization.

Additionally, a functional trans-endothelial electrical resistance (TEER) assay was performed to investigate whether the structural tight junction modifications observed herein were accompanied by alterations in endothelial monolayer barrier properties. Cultivation of endothelial cells with RG-CM for 24 h increased TEER barrier properties by 22%. However, addition of Inf-RG-CM completely impaired RG-induced increases in barrier properties, although no significant alterations were observed in cell morphology. Treatment with Inf-RG-CM decreased TEER barrier properties by 27% when compared to the controls (**Figure 2 F**).

**Figure 2.**
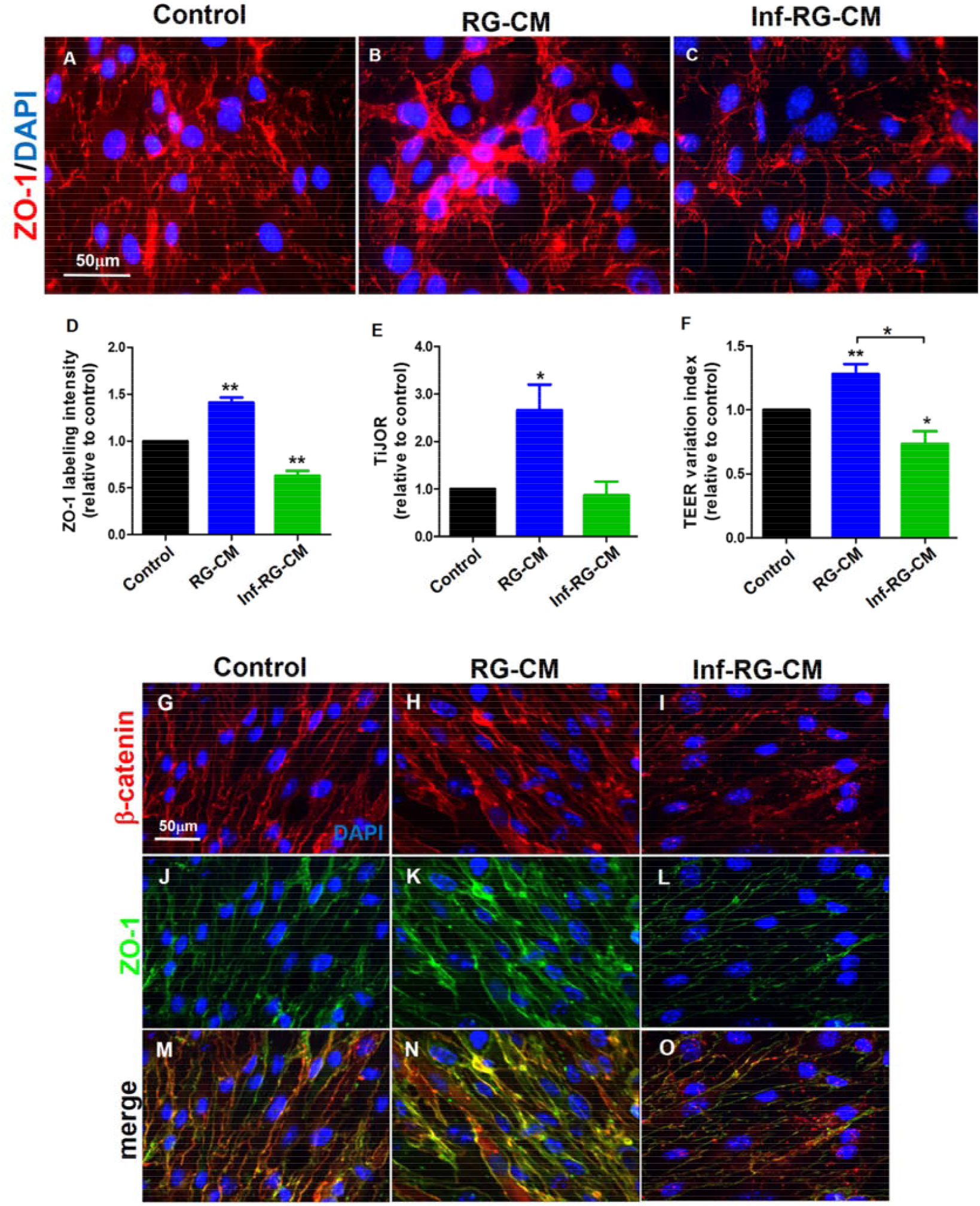
Conditioned medium from *T. gondii*-infected Radial glia decreases ZO-1/β-catenin proteins organization and impairs barrier properties of brain microvascular endothelial cells. Murine brain endothelial cells (bEnd.3) were incubated for 24h with cultivation medium (Control), conditioned medium derived from uninfected RG cultures (RG-CM) or conditioned medium derived from *T. gondii*-infected RG cultures (Inf-RG-CM). ZO-1 tight junction adapter protein was found mainly distributed along adjacent cell membranes of confluent control cultures (A). Addition of RG-CM to endothelial monolayers significantly increased the levels of ZO-1 proteins on cells surfaces (B, E). bEnd.3 cells incubated with Inf-RG-CM (C, E) presented reduced levels of ZO-1 immunoreactivity, when compared with controls (A-E). Tight junction organization index in cell-cell contact regions (TiJOR) was also increased by addition RG-CM, wheb compared with control cultures, and significantly disrupted by Inf-RG-CM presenting as discontinuous ZO-1 labeling, when compared with RG-CM-treated cultures (F). Transendothelial electrical resistance (TEER) was significantly increased by RG-CM when compared to all conditions. Inf-RG-CM treatment impaired RG potential to induce TEER by reducing electrical resistance on bEnd.3 cells bellow control levels (G). In Control cultures β-catenin was found associated with cell-cell junctional contacts, similar to ZO-1 tight junction distribution (H, K, N). Addition of RG-CM to endothelial monolayers enhanced β-catenin immunoreactivity and similar to ZO-1 protein on cells contacts regions (I, L, O). bEnd.3 cells incubated with Inf-RG-CM presented reduced immunoreactivity for β-catenin similar to ZO-1, concomitant with junctions disorganization (J, M, P) when compared with controls (A-C). β-catenin distribution in infected cells were mainly displaced to intracellular parasites *P=0.03, **P=0.0001, one-way ANOVA with Bonferroni post-test.

The role of RG infection on the distribution of the pro-angiogenic signaling protein β-catenin in endothelial cells was also investigated. The deleterious effect of Inf-RG-MC seen in TiJOR and TEER was also true for β-catenin protein distribution on bEnd.3 cells monolayers. In the control condition, β-catenin colocalized with ZO-1 along cell-cell contacts (**Figure 2 G, J, M**). The RG-CM treatment significantly increased β-catenin immunolocalization at junctional regions, which was impaired by Inf-RG-CM treatment (**Figure 2 H, K, N and I, L, O**).

### 3.3) *T. gondii-*infected RG cultures present altered cytokine secretion that affects endothelial cell barrier properties

To gain insight into the possible alterations induced by *T. gondii* on the RG secretory protein repertoire, a CBA analysis was carried out to measure the inflammation related factors IL-2, IL-4, IL-6, IL-10, IL-17, IFN-γ and TNF-α, followed by an ELISA analysis for the neuroprotector TGF-β1, to measure the levels of these cytokines in the RG-CM and Inf-RG-CM. No detectable levels of IL-2, IL-4, IL-10, IL-17, IFN-γ and TNF-α, in the conditioned mediums were observed. CM from infected RG cultures showed IL-6 in increased concentrations (7.4 pg/ml) when compared to CM from control RG-CM (0.3 pg/ml, p<0.01, unpaired *t* test). TGF-β1 levels were 40% decreased in Inf-RG-CM when compared to uninfected cultures (18 versus 29 pg/ml, p<0.05, Unpaired Student’s T test) (**Figure 3A**).

**Figure 3.**
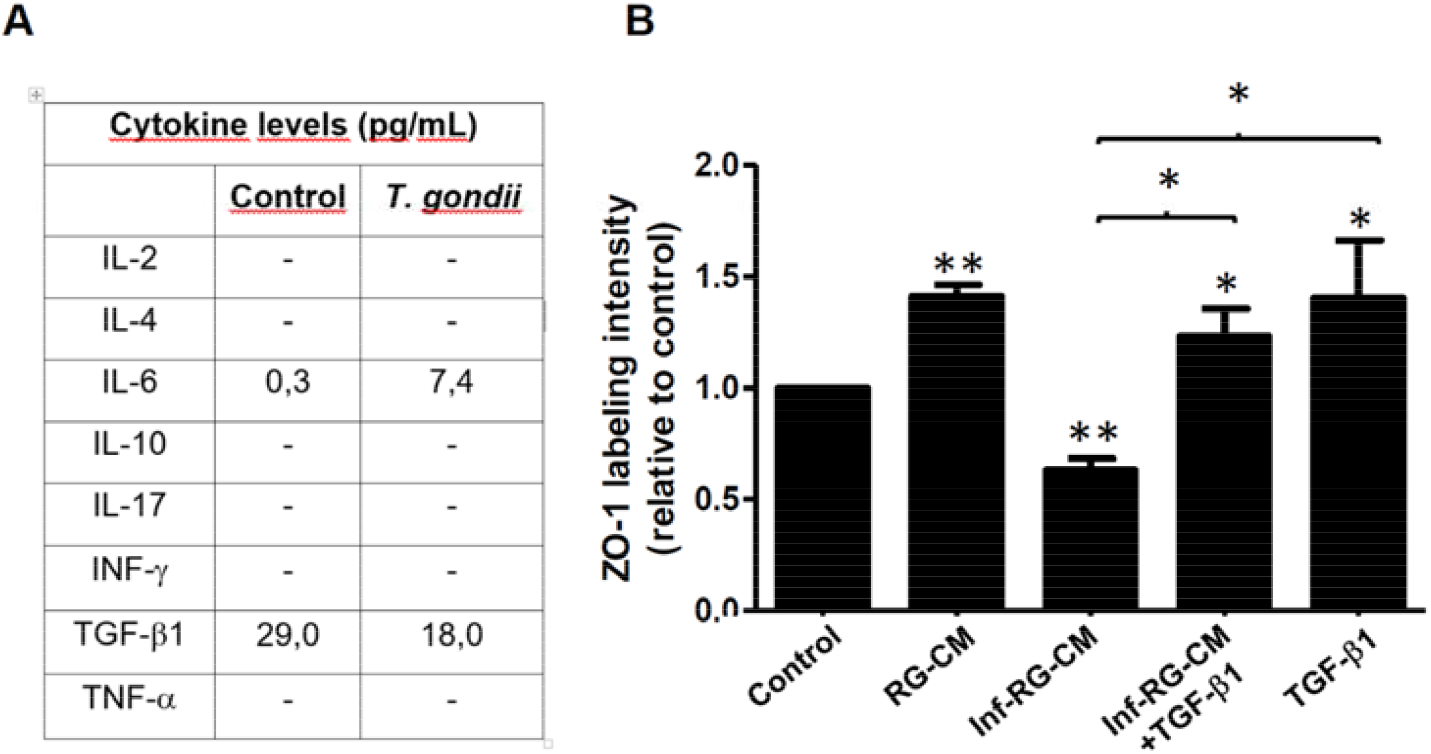
*T. gondii* infection alters the levels of secreted cytokines by Radial glia cells. Conditioned medium from uninfected RG cultures (RG-CM) and from cultures infected with *T. gondii* tachyzoites for 24 h (Inf-RG-CM), were subjected to Cytokine Bead Array (CBA) and ELISA assays to measure the levels of pro and anti-inflammatory secreted cytokines. CBA detected augmented levels of IL-6 and ELISA revealed decreased TGF-β1 levels (A) in Inf-RG-CM, compared with RG-CM. Addition of TGF-β1 (19=0ng/mL) to Inf-RG-CM significantly rescued the levels of ZO-1 labeling intensity, similar to addition of TGF-β1 alone to bEnd.3 cells, when compared to RG-CM. **p=0.0001, *p=0.036, one-way ANOVA with Bonferroni post-test.

Since TGF-β1 has a crucial role on brain microvasculature, we treated bEnd.3 cells with Inf-RG-CM together with recombinant TGF-β1 (10ng/mL) for 24 hours and performed immunocytochemistry for ZO-1. Addition of TGF-β1 to Inf-RG-CM completely rescued the ZO-1 labeling intensity to levels comparable to RG-CM condition, similarly to addition of TGF-β1 alone (**Figure 3 B**).

## 4. DISCUSSION

### 4.1) RG differentiation potential is impaired by *T. gondii* infection

RG cells have been extensively investigated concerning several features, including neuronal migration support and multipotent neural stem cell potential in generating neurons, astrocytes, oligodendrocytes and other progenitor subtypes in the cerebral cortex (Rakic, 1999; Noctor et al., 2001; Morest and Silver, 2003; Barnabe-Heider et al., 2005; Stipursky and Gomes, 2007; Kessaris et al., 2008; Kriegstein and Alvarez-Buylla, 2009; Ortega and Alcantara, 2010; Stipursky et al., 2012; Stipursky et al., 2014). Evidence suggest that neural progenitors are directly affected by the TORCH complex of perinatal infectious diseases that, ultimately, lead to malformations in the cerebral cortex, such as microcephaly, mostly by disrupting neural cells generation (Neu et al., 2015). Regarding infection with *T. gondii*, recent findings point to increased apoptosis and reduced cell differentiation in the C17.2 neural stem cell line (Gan et al., 2016; Wang et al., 2014; Zhang et al., 2017). However, further characterization and evaluation of the effects of *T. gondii* infection on primary RG cells isolated from embryonic cerebral cortex have not yet been addressed. The present study indicates that *T. gondii* significantly decreases the number of Nestin/BLBP positive RG cells, possibly by decreasing their proliferation. Accordingly, previous data describe that altered neural stem cell proliferation, differentiation and apoptosis can be triggered by viral infection (Gan, 2016; Souza et al., 2016). In our model, RG cells accounted for 51% of the total cell population in the control condition after a total period of 48 h of cultivation. Since the total number of cells and the small percentage of apoptotic RG cells were not affected by *T. gondii* infection, it is possible that global cytotoxic infection effects were not the mechanism leading to reduced numbers of, specifically, Nestin/BLBP positive RG cells. Although RG cells have been described by our group and others to differentiate into astrocytes at later stages of cortical development (Rakic, 1971; de Azevedo et al., 2005; Stipursky and Gomes, 2007; Stipursky et al., 2012; Stipursky et al., 2014), no alterations in astrocyte differentiation by *T. gondii* infection were observed. Secreted levels of IL-6 and TGF-β1, two cytokines known to positively mediate astrocyte differentiation from neural progenitor cells (Taga and Fukuda, 2005; Nakamura et al., 2005, Stipursky and Gomes, 2007; Stipursky et al., 2012; Stipursky et al., 2014), were altered in Inf-RG-CM. However, it is possible that, in this context, altered cytokine levels might not control autocrine regulation of astrocytogenesis, or that other molecular mechanisms known to modulate gliogenesis are not altered in this context. On the other hand, a reduction in β-III-tubulin positive cell numbers was detected, without affecting apoptotic neuronal population, suggesting inhibition of neurogenesis. This finding is corroborated by the recent demonstration that *T. gondii* infection impairs neuron generation from C17.2 neural stem cell line *in vitro* (Gan et al., 2016; Zhang et al., 2017). However, it is possible that, in addition to *T. gondii* induced alterations of well-known molecular mechanisms that control neurogenesis during cortical development, such as Wnt/βcat signaling (Gan et al., 2016; Zhang et al., 2017) and neuronal differentiation transcription factors, such as Fabp7/BLBP, Sox2, Tacc3 Eya1, Sox2, and Tnfrsf12a genes (Xiao et al., 2012), altered levels of IL-6, TGF-β1 and other, still unidentified cytokines, might exert an autocrine effect on the neurogenic potential of RG cells. Although *T. gondii* infection decreased the number of Nestin/BLBP and β-III-tubulin positive cells and did not alter GFAP cells numbers, overall cell composition percentages induced an increase in an unidentified cell population, suggesting that RG differentiation might favor the appearance of other cell types. Thus, since the literature lacks a more detailed investigation regarding the impact of congenital transmission of *T. gondii* in cerebral cortex embryonic development *in vivo*, it is essential to address how neurogenesis and/or gliogenesis are affected during CT in the developing brain.

### 4.2) Impacts of *T. gondii* infection on RG potential to control endothelial barrier properties

Vascular development by angiogenesis, in addition to blood vessel stability and function, result from a fine-tuned control of pro- and anti-angiogenic molecules produced by endothelial and neighboring cells, as well as environmental cues (Carmeliet and Jain, 2000). In the last few years, RG have been pointed out as an essential cellular and molecular scaffold for blood vessel formation and vascular stability acquisition during cerebral cortex development (Ma et al., 2013; Errede et al., 2014; Hirota et al., 2015; Siqueira et al., 2017). Herein, we demonstrate that RG-CM treatment of endothelial cells greatly increases tight junction ZO-1 protein levels and organization, suggesting that RG-secreted factors promote microvascular barrier formation. Vascular stability and barrier properties are essential features that allow the controlled transport of nutrients and other substances across the BBB (Abbott, 2005; Ben-Zvi et al., 2014). Several molecular mechanisms were shown to promote the expression of tight junction proteins Claudin-5, Occludin and ZO-1 in endothelial cells, thus leading to the formation of the BBB, including activation of the TGF-β1 signaling pathway, PDGFR/PDGR-B interaction and Wnt/βcatenin (Alvarez et al., 2011; Baeten and Akassoglou, 2011; Zhao et al., 2014; Zhao et al., 2015). In this context, *T. gondii* infection may indirectly deregulate signaling pathways critical in controlling vessel stability by (i) affecting RG potential concerning mediation of BBB formation or (ii) disrupting tight junction proteins expression and organization directly in endothelial cells. Increased BBB disruption and permeability have been recently correlated t elevated levels of the pro-inflammatory cytokine IL-6 in the cerebrospinal fluid of human adult individuals affected by neuromyelitis optica and neuropsychiatric systemic lupus erythematosus, and in a rat model of psychosocial stress induction (Uchida et al., 2017; Asano et al., 2017; Schiavone et al., 2017).

Pro-inflammatory cytokines, such as IL-6, have been previously reported to promote BBB dysfunction in several neurodegenerative disease contexts, such as Alzheimer’s, Parkinson’s and Multiple sclerosis, as well as CNS ischemia and infectious diseases, being a potential neuroinflammation establishment mechanism (Kempuraj et al., 2016; Rochfort et al., 2016). Stimulation of endothelial human brain cells (HCECs) with IL-6 was shown to induce VEGF synthesis and release, promoting angiogenesis, alongside increased expression of matrix metalloproteinase 9 (MMP9) (Yao et al., 2006). Increased MMPs levels in different brain injury models were associated with degradation of the endothelial basement membrane and enhanced tyrosine phosphorylation of tight junction proteins, triggering protein redistribution, changes in adhesive properties between endothelial cells, and TEER reduction, resulting in increased permeability (Stamatovic et al., 2008).

Although the roles of IL-6 have been pointed as essential to endothelial function, angiogenesis and blood vessel stability have been classically described as depending on anti-inflammatory TGF-β1 cytokine signaling in the embryonic and adult brain (Dohgu et al., 2004; Lebrin et al., 2005; Holderfield and Hughes, 2008; Arnold et al., 2014; Hellbach et al., 2014; Siqueira et al., 2017). TGF-β1 is a multifunctional cytokine that controls multiple physiological and pathological events, such as embryogenesis, immune response, ECM synthesis, cell differentiation and cell-cycle control in several tissues (Massague, 1998; Massague and Gomis, 2006). TGF-β1 in the CNS has been reported to play a key role in neuronal generation, survival and migration (Brionne et al., 2003; Miller, 2003; Esposito et al., 2005; Stipursky et al., 2012), glial differentiation (Sousa Vde et al., 2004; Romao et al., 2008; Stipursky et al., 2014), and synapse formation (Diniz et al., 2012; Diniz et al., 2014). More recently, our group demonstrated that TGF-β1 secreted from RG cells induces angiogenesis in the developing cerebral cortex (Siqueira et al., 2017). Herein, we observed that CM from infected RG cells contains lower TGF-β1 levels compared to uninfected cells. Loss of the active form of TGF-β1 cytokine, mutation of the *Tgfb1* gene or even deletion of *Tgfr2* or *Alk5/Tgfbr1* genes in endothelial cells of the embryonic forebrain have been shown to promote excessive vascular sprouting, branching and induce cerebral hemorrhage (Arnold et al., 2014). Furthermore, TGF-β1 is known to promote tight junction proteins and P-glycoprotein transporter expression in brain endothelial cells (Dohgu et al., 2004), induce endothelial barrier properties, such as γ-glutamyl-transferase (GGT) expression, mediated by astrocyte secretion (Garcia et al., 2004), and TGF-β signaling is involved in later stages of blood vessel development, such as the induction of maturation and stability maintenance mediating the interaction between endothelial and mural cells (Lebrin et al., 2005).

Here we demonstrated that Inf-RG-CM presented increased amounts of IL-6 and significantly less TGF-β1, compared to control RG-CM. Evidence shows that disruption of endothelial interactions with neurovascular unit or low levels of TGF-β1 leads to abnormal distribution of junctional proteins and increased vascular permeability (Garcia et al., 2004; Dohgu et al., 2005; Winkler et al., 2011), In the BBB, TGF-β1 secretion by astrocytes has been demonstrated as essential for maintaining brain vasculature stability and inhibiting or decreasing leukocyte transmigration across the endothelium (Fabry et al., 1995; Alvarez et al., 2013). Since mature BBB astrocytes are the main source of TGF-β1, and RG cells also secrete TGF-β1, which we demonstrated as promoting RG-astrocyte differentiation (Stipursky and Gomes, 2007; Stipursky et al., 2012, 2014), it is possible that, in our infection model, reduced levels of RG-derived TGF-β1, and possibly elevated levels of IL-6 proinflammatory cytokine, might impair endothelial ZO-1 tight junction organization and β-catenin association in adherens junctions.

By adding TGF-β1 to the Inf-RG-CM, there was a complete rescue of endothelial ZO-1 protein levels, suggesting that *T. gondii* infection impairs TGF-β1 expression/secretion, or both, by RG cells.

In this context, alteration of RG secretome by *T. gondii* infection, may act as underlying mechanisms of disruption of endothelial cell barrier properties. However, specific downstream angiogenesis and vascular maturation molecular targets have not yet been identified.

Herein, β-catenin has been pointed out as a potential target for RG-endothelial dysfunctional interaction. The Wnt/β-catenin pathway has been extensively investigated and demonstrated as exherting critical roles on vascular development and maturation in the developing cerebral cortex and maintenance of BBB in adult brain (Liebner et al., 2008; Daneman et al., 2009; Ma et al., 2013; Engelhardt and Liebner, 2014). β-catenin, the canonical downstream mediator of Wnt signaling, associates with classic cadherins in adherens junctions and to α-catenin, that mediates its association with the actin cytoskeleton. In endothelial cells, β-catenin is constitutively bound to VE-cadherin and when β-catenin is free in cytoplasm, it is rapidly inactivated by phosphorylation and ubiquitination by a protein complex that includes axin and adenomatous polyposis coli (APC). Upon downregulation of VE-cadherin expression or Wnt signaling activation in endothelial cells, β-catenin translocates to the nucleus to modulate gene expression, acting as a cotranscription factor together with lymphoid enhancer factor/T-cell factor (Lef/TCF), Forkhead box protein O1 (FoxO1), hypoxia-inducible factor (HIF), Smads and others, mediating cellular responses such as cell cycle, apoptosis, cell differentiation and cell-cell communication (Klaus and Birchmeier, 2008; Giannotta et al., 2013).

A previous report demonstrated that VE-cadherin phosphorylation prevents p120 and β-catenin binding, triggering destabilization of adherens junctions, maintaining cells in a mesenchymal state (Potter et al., 2005).

Endothelial cells adherens and tight junctions lay in close proximity to neighbor cells, both mediating membrane adhesion and limiting paracellular permeability. Although adherens and tight junctions are formed by distinct structural proteins, with specific binding affinities, the assembly of tight junctions has been demonstrated as coupled with the formation of adherens junctions. Physical linkages between these junctional structures can be mediated by a multistep process that involves α–, β-catenins, ZO-1 and afadin proteins interactions, initiating with adherens junctions protein clustering and structural organization on cell membranes that later recruit and allow tight junctional machinery to assemble (for revision Campbell et al., 2017). In fact, β-catenin/FoxO1 transcription factor complex was demonstrated to repress Claudin-5 tight junction gene expression (Taddei et al., 2008). Ma and colleagues (2013) demonstrated that Wnt/β-catenin signaling activation can mediate RG-endothelial interaction through transcriptionally induction of MMP2/MMP9 in endothelial cells, which was correlated to decreased vessel stability and increased endothelial proliferation in the developing cerebral cortex. Accordingly, increased MMPs levels were found in astrocyte cultures (Lu and Lai, 2013) and in the sera of pregnant women infected with *T. gondii* (Wang and Lai, 2013), suggesting that Wnt/β-catenin signaling may modulate junction protein levels expression and organization and/or turnover though multiple mechanisms. Thus, in our context, *T. gondii* infection effects might exert critical function in triggering endothelial junction dismantling, alongside β-catenin dissociation from cell-cell contacts.

Together, our results suggest that *T. gondii* deregulates RG cells proliferation, neurogenesis potential and secretory profile, with decreased TGF-β1 and, to a less extent, increase in IL-6 levels. In addition, the potential of RG cells to modulate endothelial cell function is also affected by *T. gondii* infection, resulting in deficient organization of tight junction protein ZO-1 and junction associated β-catenin, reduced TEER, leading to impaired endothelial stabilization and loss of barrier properties. In a CT context, alterations in the RG cell differentiation potential and in RG-endothelial cell interactions may be critical to the reduced numbers of neurons generated during cortical development and dysfunctional BBB formation, which would directly contribute to the establishment of the microcephaly phenotype. Although RG-endothelial interactions are critical to promote vascular development and BBB formation, the specific molecular mechanisms disrupted by *T. gondii* infection in such interactions are not known. Thus, an improved description of the signaling pathways involved in such events might contribute to the development of therapeutic approaches to rescue or protect neural stem cells functions and vascular development, thus preventing the clinical manifestations observed in CT.

## 5) ACKNOWLEDGEMENTS

We thank Dr. Flávia Carvalho Alcantara Gomes for providing equipment, laboratory facility and some reagents; Marcelo Meloni, Adiel Batista do Nascimento and Sandra Maria Oliveira Souza for technical assistance and Dr. José Morgado Diaz (INCA) and Dr. Luzia Maria de Oliveira Pinto (IOC, Fiocruz) for the use of their MilliCell equipment.

## 6) AUTHOR CONTRIBUTIONS STATEMENT

<u>DA</u> performed bEnd.3 cultures, infection and TEER experiments. <u>ACM</u> performed bEnd.3, treatments, immunocytochemistry quantifications and radial glia immunocytochemistry quantifications. <u>MS</u> performed radial glia and bEnd.3 immunocytochemistry. <u>CMC</u> and <u>MCW</u> performed CBA and ELISA cytokines analysis, respectively. <u>JS</u> performed radial glia cultures, bEnd.3 immunocytochemistry, TGF-β1 ELISA and wrote the first draft of the manuscript. <u>HSB</u> discussed the experimental design and data interpretation regarding *T. gondii* infection, provided equipment, laboratory facility and some reagents. <u>DA</u> and <u>JS</u> equally contributed in the design of most of the experiments. All authors contributed to manuscript revision, read and approved the submitted version.

---

**Supplementary figure 1.**
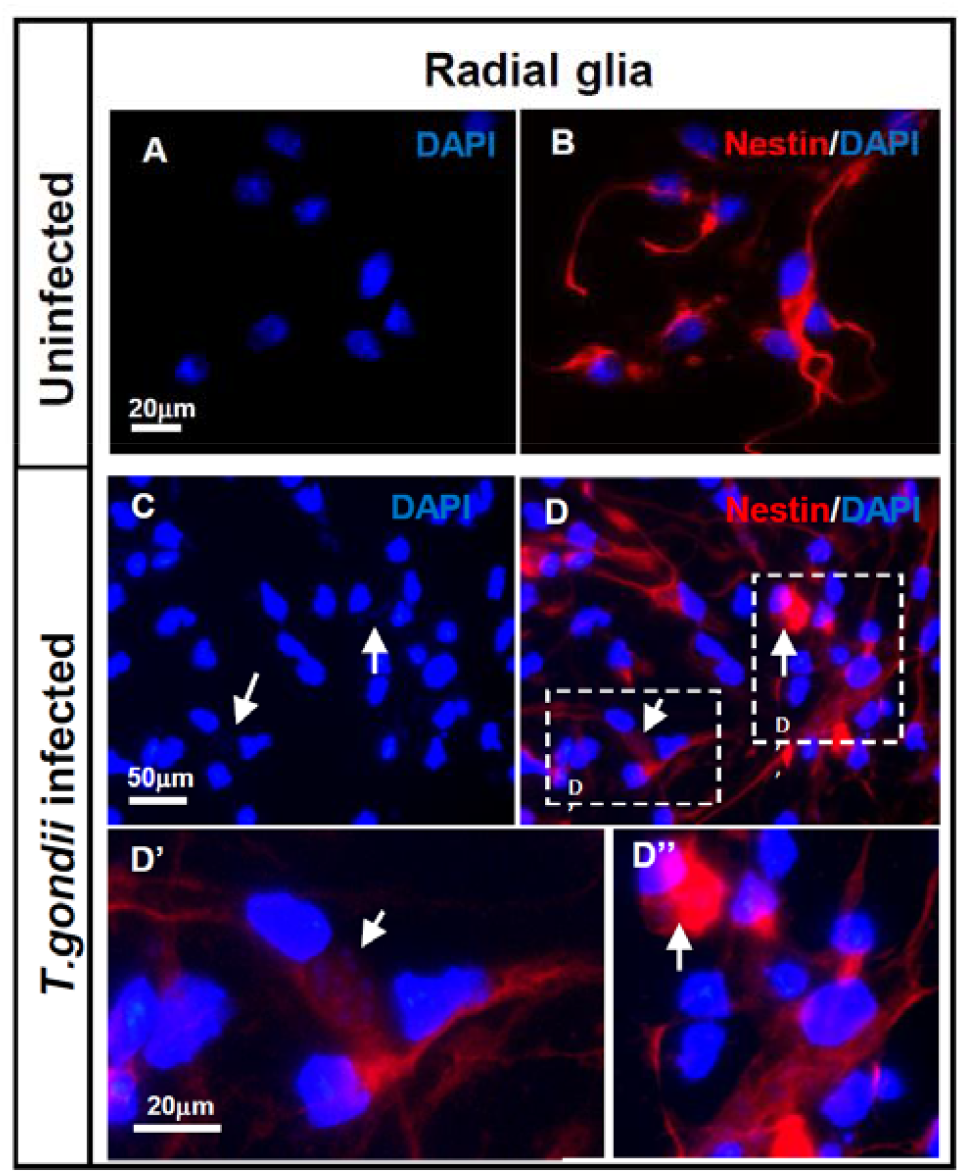
Identification of *T. gondii* tachyzoites in infected cells. RG cultures were infected with *T. gondii* tachyzoites for 24h and labelled with DAPI to identify host cell nucleus and parasite DNA (blue). Uninfected RG cultures labeled with DAPI and Nestin (red) (A, B). Infected RG cells showing host and tachyzoites DNA in the cytoplasm (C, D, arrows), high magnification of tachyzoites in infected cells (D’, D”).

**Supplementary figure 2.**
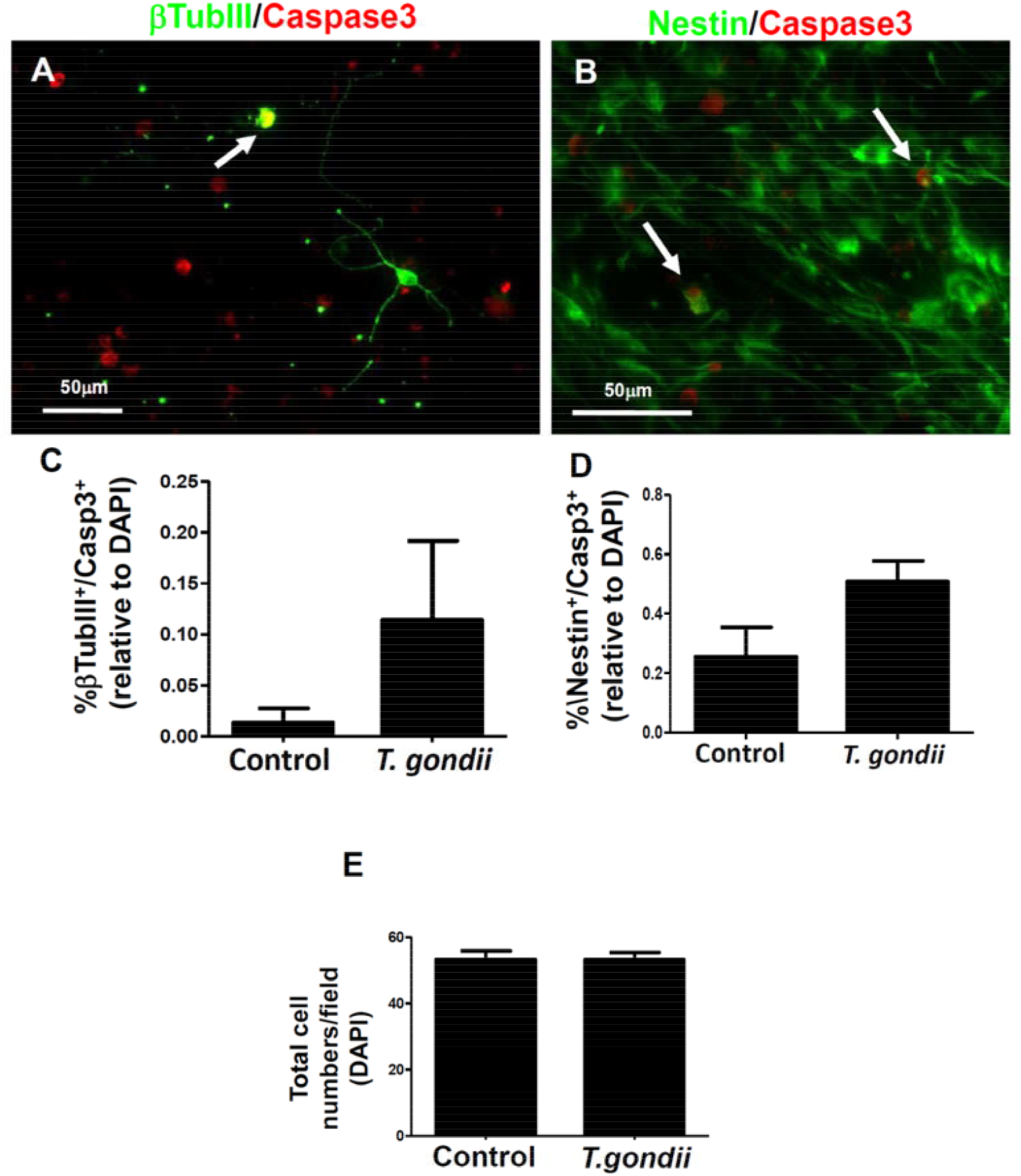
*T. gondii* infection does not affect RG and neuronal cell death. RG cultures were infected with *T. gondii* tachyzoites for 2h and and labelled with neuronal marker (bTubulin), RG marker (Nestin) and apoptotic cell marker (cleaved-Caspase 3), No statistic differences in the percentage of β-tubulinIII/Caspase3 (A, C) and Nestin/Caspase3 (B, D) double positive cells were observed. Quantification of total DAPI-labeled nucleus per microscopical field in uninfected and infected RG cultures revealed no significant difference in total cells number in both conditions (E). P>0.05, unpaired t-test.

